# Deep Learning-Based Inference of Uterine Contractions from Maternal ECG

**DOI:** 10.1101/2024.09.29.615726

**Authors:** Shunsuke Tanaka, Keisuke Ito, Kyohei Takano, Yohsuke Takasaki

**Affiliations:** nonat Inc., Tokyo, Japan; Saintpaulia Medical Corporation Misao Ladies’ Hospital, Gifu, Japan; Okayama University, Okayama, Japan; University of Cambridge, Cambridge, United Kingdom; Hiroshima University, Hiroshima, Japan

## Abstract

**Background:** Preterm birth and stillbirth have multifaceted causes, many of which remain unknown. Factors such as maternal age, sexually transmitted infections, and genetic abnormalities are notably implicated. Globally, there has been minimal improvement in preterm birth rates, leading to substantial economic losses and underscoring the need for innovative approaches.

**Objective:** This study aims to analyze and detect uterine contractions by applying deep learning techniques to physiological data derived from maternal electrocardiograms (ECGs). Leveraging ECGs, which are easily obtainable from wearable devices, allows for effective and convenient monitoring of uterine contractions outside clinical settings. Since uterine contractions are a critical indicator for assessing the risk of preterm birth, our algorithm has the potential to facilitate early identification of high-risk individuals, thereby contributing to timely interventions and improved maternal-fetal outcomes.

**Methods:** Participants meeting all inclusion criteria and none of the exclusion criteria were recruited from patients admitted to or attending outpatient services at Saintpaulia Misao Ladies Hospital in Gifu, Japan, between December 6, 2023, and July 31, 2024. Our deep learning model was developed using maternal ECG data, uterine contraction waveforms obtained from cardiotocograms (CTGs). The collected data were divided into training and evaluation datasets. The AI-generated uterine contraction waveforms from the developed model were compared with ground truth labels obtained from the fetal monitoring devices. This study was reviewed and approved by the Ethics Committee of the Graduate School of Medicine at Gifu University and was conducted as a collaborative research effort with nonat Inc. (the lead research institution) and Saintpaulia Misao Ladies Hospital (joint research institution).

**Results:** Finally, 73 participants took part in this study, and 57 datasets were used for algorithm development and evaluation. Multiple measurements were taken on the same subjects on different days only with their consent. Our analysis demonstrated a strong correlation between the uterine contraction waveforms estimated by the developed model and the ground truth waveforms from CTGs, achieving an average correlation across all evaluation data (Pearson correlation coefficient = 0.53).

**Conclusion:** This is the world’s first trial to detect uterine contractions from maternal ECGs using deep learning algorithms. In this study, we successfully developed a deep learning algorithm capable of accurately inferring uterine contraction waveforms from maternal ECGs. Given that ECGs can be easily obtained using wearable devices, this approach may provide healthcare professionals with objective and precise information on uterine contractions—a crucial indicator of preterm labor—even in resource-limited settings outside hospital environments.

## Introduction

### Background on Uterine Contractions and Preterm Birth

Uterine contractions are a critical clinical indicator in perinatal care(1), essential for predicting childbirth and assessing the risk of threatened preterm birth (2). However, accurately evaluating uterine contractions has long been a challenge in the field of perinatal medicine (3). A study has shown that only about 10% of high-risk preterm birth patients, diagnosed with high-risk uterine contractions, actually delivered within a week, indicating that unclear interpretations of uterine contractions may lead to unnecessary treatments (4).

Preterm birth itself is a significant social issue in perinatal care (5). While various factors, such as maternal age, infections, and genetic abnormalities, have been identified as potential causes, many remain unknown (6). The risk of preterm births is also rising due to changes in social factors, including the trend toward advanced maternal age (7)(8)(9)(10)(11). A 2020 WHO report highlighted the global burden of preterm births, describing it as a “silent emergency” due to the lack of improvement in preterm birth rates over recent decades (5).This situation underscores the limitations of modern medicine in addressing threatened preterm birth and suggests the presence of unmet needs in this area (12).

### Medium- to Long-Term Risks for Preterm Infants

Preterm births and stillbirths are not merely individual concerns; their societal impact is profound (13)(14). The extensive costs associated with tertiary care and ongoing life support are significant (14). In the United States, over $20 billion is spent annually on managing preterm births (14), while in the United Kingdom, nearly £2.5 billion is allocated to address this issue, underscoring the considerable societal burden of preterm births(15).

In addition, patients born preterm face increased risks of various complications. Specifically, these infants are more susceptible to hypothermia, hypoglycemia, respiratory distress, apnea, hyperbilirubinemia, feeding difficulties, low Apgar scores (below 4), and neurological issues such as seizures and perinatal asphyxia (16)(17)(18)(19)(20)(21). Even beyond the neonatal period, preterm infants are at higher risk for long-term complications, including cognitive impairment and lower academic achievement, which are neurodevelopmental outcomes that often manifest in childhood or adulthood (22)(19).

### Significance of This Study

This study is a physiological and technical exploratory research based on the hypothesis that there is a mapping relationship between the electrical activity recorded from the maternal body surface and the characteristics of fetal heart rate and uterine contractions (intensity, duration, and interval), and that this relationship can be inferred through features learned by deep learning. Currently, in the medical field, there are numerous examples where deep learning is utilized in clinical practice (23)(24)(25)(26). In particular, the application of deep learning to time-series data (e.g., LSTM or Transformers) has the potential to become a powerful tool that could replace traditional group-to-group comparisons (27).

The purpose of this study is to support medical professionals in identifying high-risk preterm birth patients in out-of-hospital settings, where medical resources are limited. Given that ECGs are easily available from wearable devices, leveraging this approach allows uterine contractions to be monitored effectively and conveniently outside clinical environments. By utilizing this easily obtainable biometric information, our research aims to illuminate the “black box” of pregnancy status outside hospital environments and help medical professionals more effectively detect those who are at high risk of preterm birth. This may potentially reduce unnecessary medical interventions and contribute to timely interventions, leading to improved perinatal outcomes.

## Method

This research plan has been reviewed and approved by the Ethics Review Committee of the Graduate School of Medicine at Gifu University, and the study has been conducted with the permission of the heads of each collaborating research institution.

Participants who provided informed consent for this clinical study and met the inclusion criteria were equipped with ECG measurement sensors during Non-Stress Test (NST) or Cardiotocography (CTG) examinations using a fetal monitoring device. The ECG signals were recorded at a sampling rate of 1000 Hz for a duration of 15 to 40 minutes, capturing Lead I.

The inclusion criteria were as follows:

- Individuals who fully understood the research plan and were capable of providing written informed consent.
- Healthy adult pregnant women at 37 weeks of gestation or later.
- Singleton pregnancy.

The exclusion criteria were as follows:

- Pregnant women with obstetric complications.

### Collected Data

- **Test Data**: Ultrasound, cardiotocography (CTG), electrocardiography (ECG)
- **Medical Records**: Age, height, weight, blood pressure, vital signs, urinalysis results (urinary glucose, urinary protein), medication history, obstetric history (gravida and para), gestational age, ultrasound findings, delivery outcomes (mode of delivery, obstetric complications, infant birth weight, Apgar score, umbilical arterial blood pH), pregnancy background (natural conception, artificial insemination, assisted reproductive technology (ART), pregnancy outcomes (miscarriage, ectopic pregnancy, heterotopic pregnancy), induced abortion, live birth, stillbirth, history of fetal reduction surgery, number of births), and pre-pregnancy medical history.

## Data processing

In this study, data preprocessing was performed to enable efficient analysis of long-term ECG data by reducing its size. Instead of directly inputting raw data obtained from biometric monitors into the model, we conducted feature extraction of the ECG signals. Specifically, we utilized an ECG waveform feature extraction library, NeuroKit2, to automatically classify the ECG waveforms and identify the key waveform components of the ECGs, such as the P wave, Q wave, R wave, S wave, and the time points and voltage values of the baseline peaks.

The rationale behind this preprocessing was that simple downsampling might risk losing crucial information relevant to inferring uterine contractions, and thus, meaningful data reduction was necessary. Furthermore, as shown in Figure 1, distinguishing between contraction and non-contraction periods in uterine contraction waveforms is challenging. Therefore, we created annotated data that reflected expert opinions from obstetricians.

**Figure 1.**
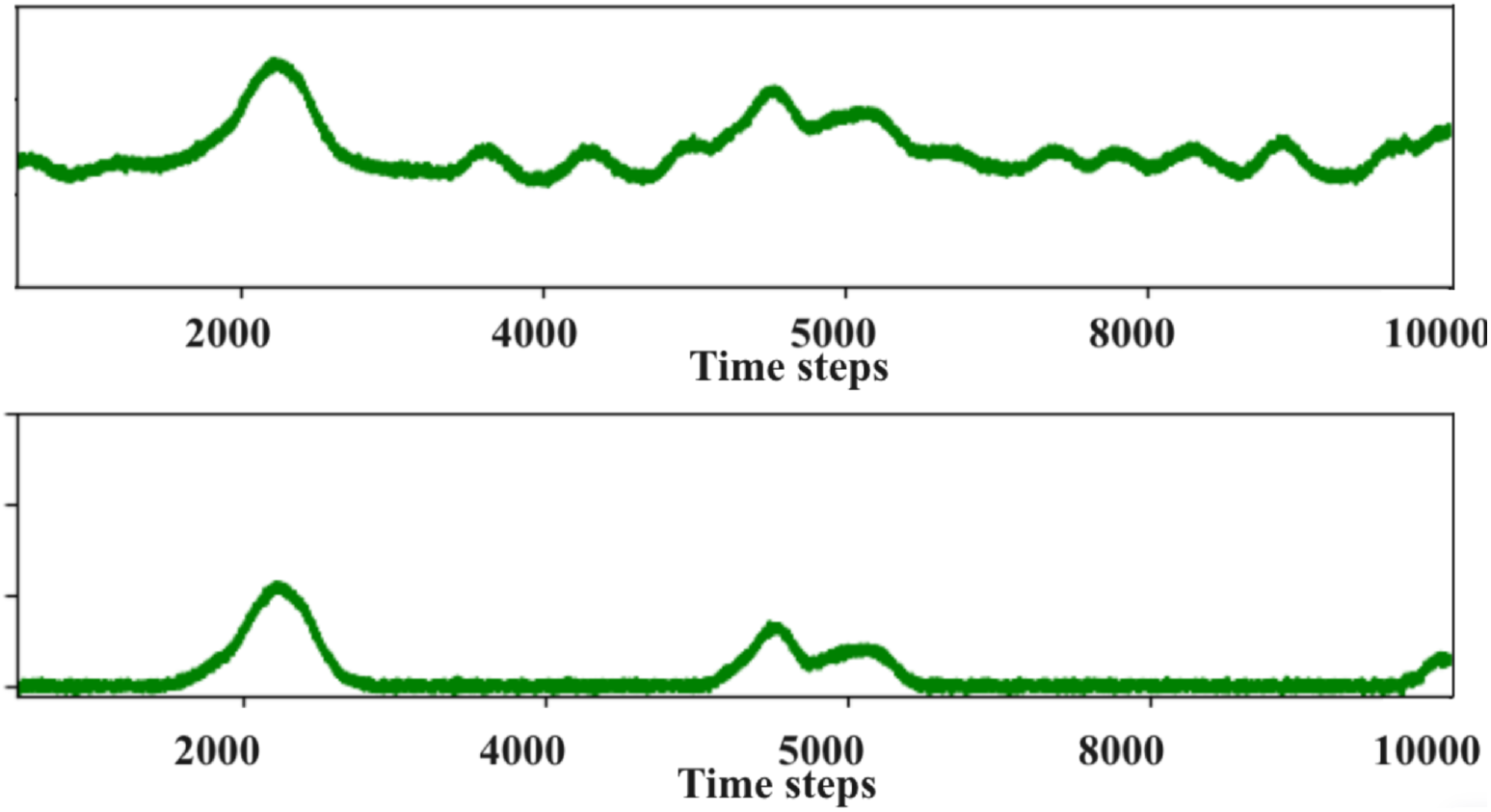
Annotated Diagram of Uterine Contractions During Contraction and Non-Contraction Periods

To facilitate the annotation process, we developed software that makes it easier to distinguish between contraction and non-contraction periods. This software displays the uterine contraction waveform on the screen, allowing physicians to click on the start and end points of the contraction periods, which records the time, while adjusting the y-coordinates of non-contraction periods to align with the baseline. We compared two types of data: one that used the annotated data as ground truth labels for training and the other using the original unannotated data, to evaluate the performance.

## Development of AI model optimized Long-term series

In this study, we constructed a sequence-to-sequence Long Short-Term Memory (LSTM) model (Figure 2) to predict one time-series data (uterine contractions) from another time-series data (ECG). To handle long-term time-series data, we propose a novel approach that overcomes the limitations of conventional LSTM models. This is an unprecedented approach to this specific dataset, and we referred to time-series prediction models such as ECG atrial fibrillation (AF) detection and fetal heart rate prediction using LSTM (28)(29)(30)(31).

**Figure 2.**
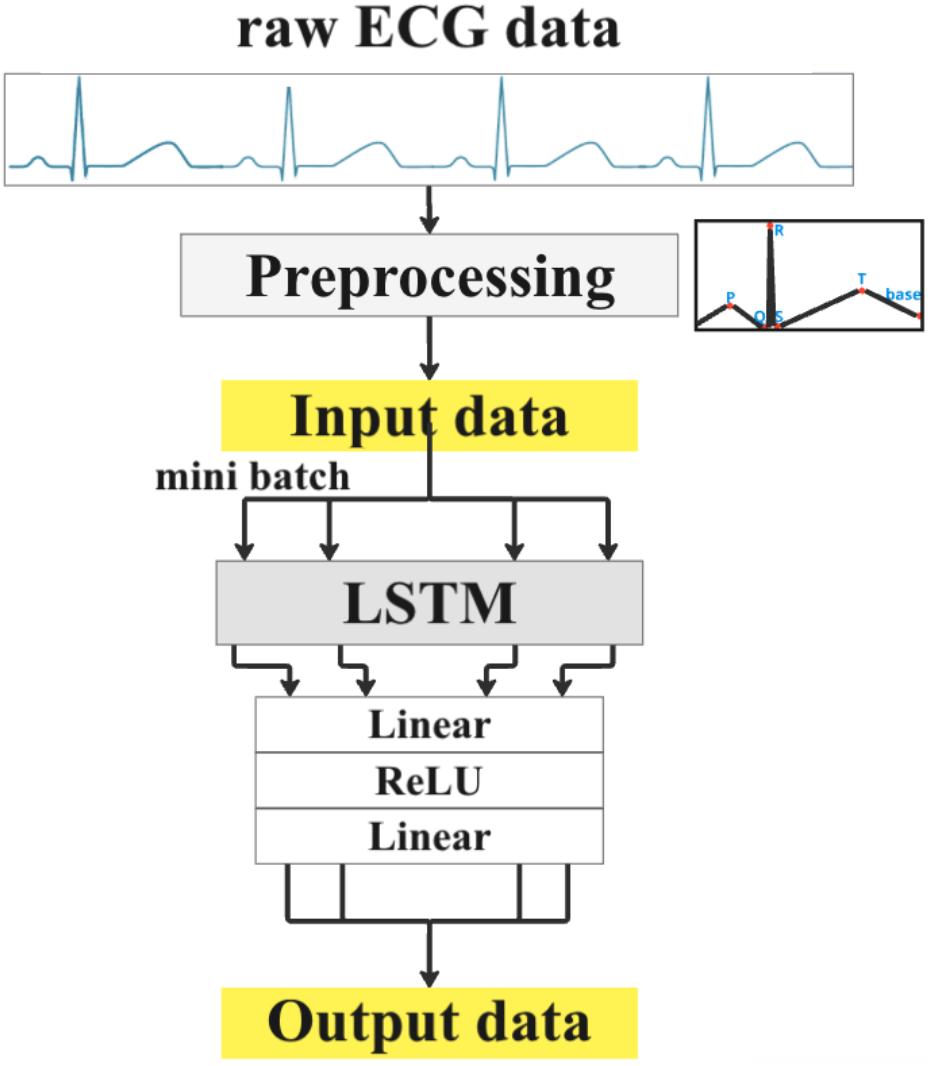
The constructed LSTM model

There are two main reasons why we chose LSTM for this study. First, LSTM is well-suited for processing large datasets. Uterine contractions involve long-term variations, requiring extensive data for analysis. Models using attention mechanisms scale computational complexity quadratically with respect to input length, whereas LSTM scales linearly, making it easier to design the model. Additionally, Transformer models have more parameters than LSTMs. While models with a large number of parameters have higher representational power, they also carry a higher risk of overfitting. Since this is a technical exploratory study with a limited amount of data, we opted for LSTM, which operates with relatively fewer parameters to avoid the risk of reduced generalization performance.

Second, uterine contractions are often described as periodic events, where a previous contraction may influence subsequent contractions. LSTM’s memory cell architecture is well-suited for capturing this type of temporal relationship in time-series data. Based on these considerations, we determined that LSTM was the optimal model for this task.

Traditional LSTMs are known to face issues such as vanishing gradients and memory constraints when learning long-term dependencies. To address these challenges, we implemented techniques to truncate backpropagation at specific points and divided the measurement data into smaller mini-batches for model training (32). The enhanced LSTM was able to retain long-term information while preserving the intrinsic characteristics of the hidden states and memory cells. This allowed for consistent information to be utilized across time, thus improving the efficiency of the learning process.

### Loss Function

The model was evaluated using Pearson correlation coefficient. Pearson correlation coefficient can assess the correlation between two datasets and is calculated independently of the scale of the data, making it a suitable metric for capturing the trends between the two datasets.

The non-stress test (NST), which is used as the ground truth label, is measured by attaching sensors to the surface of the abdomen. Factors such as abdominal fat thickness, abdominal shape, and changes in body position can affect the sensitivity of the measurements. Due to these factors, the waveform height obtained from the NST does not necessarily reflect the intrauterine pressure accurately(33). Considering that the NST waveforms capture the overall pattern of uterine contractions rather than precise intrauterine pressure, we concluded that evaluating the model using Pearson correlation coefficient was appropriate.

Pearson correlation coefficient is sometimes used for evaluating biometric waveform data. Typically, it is calculated over predefined intervals and is often applied to evaluate waveforms obtained from ECGs and PPG sensors (34)(35). It is known that Pearson correlation coefficient tends to fluctuate based on the sample size of the data. When the sample size is large, noise within the data tends to average out, resulting in a higher correlation coefficient that reflects the overall trend. Conversely, when the sample size is small, noise has a relatively larger influence, potentially lowering the correlation coefficient. Taking these factors into consideration, we carefully examined the optimal sample size for this study.

Clinically, non-stress tests are typically measured and evaluated over a 20-40 minute period. On the other hand, in medical practice, labor contractions are defined as painful uterine contractions occurring at 10-minute intervals. Based on this definition, we determined that an evaluation window of 10-20 minutes would be necessary for an effective assessment of uterine contractions in this study. Particularly, evaluations of less than 20 minutes are more susceptible to the influence of noise in the data. In this challenging context, we carefully explored appropriate models for evaluation.

Since the data length was not consistent across subjects, we employed mini-batch processing to standardize the input size and address this issue. Specifically, in this study, we set the mini-batch length to approximately 15 minutes for the evaluation.

## Results

### Descriptive Statistics of Study Participants

Biometric data was collected from subjects who met the eligibility criteria and consented to participate in this study, and the data was then divided into training and evaluation datasets for analysis. For data that was recorded for more than 25 minutes, segmentation was performed, and the data was distributed between the training and evaluation datasets. Ultimately, as shown in this table 1, data from 57 subjects were analyzed, with the training dataset having an average age of 32.2, gestational weeks of 31.4, and an average measurement time of 22.8, and the evaluation dataset having an average age of 31.4, gestational weeks of 39.1, and an average measurement time of 12.4. The split of training:validation is 7:3.

**Table 1:** Descriptive statistics of the participants’ data for both training and validation sets.

|  | Train Data |  | Validation Data |  |
| --- | --- | --- | --- | --- |
|  | Mean | Standard deviation | Mean | Standard deviation |
| age | 32.2 | 4.1 | 31.4 | 4.6 |
| mother_weight[kg] | 62.73 | 8.77 | 65.05 | 9.55 |
| mother_height[cm] | 157.5 | 5.82 | 158.1 | 5.85 |
| mother_BMI | 25.28 | 3.03 | 26.04 | 3.52 |
| days | 273.8 | 7.97 | 273.7 | 8.18 |
| fetus_weight[g] | 3084.6 | 407.69 | 3120.1 | 432.30 |

Data that was deemed difficult to analyze due to reasons such as ECG noise caused by maternal body movement and poor display of uterine contraction waveform due to improper monitor attachment was excluded from the analysis.

### Compare models

As shown in Table 2, multiple models were examined. First, we explored loss functions by evaluating model A and model C. The models were trained using either MSE or Pearson correlation coefficient in a predefined mini-batch size. Next, the models were trained using the annotated data, where the uterine contraction waveforms—the ground truth labels—had been annotated as described earlier.

**Table 2:**
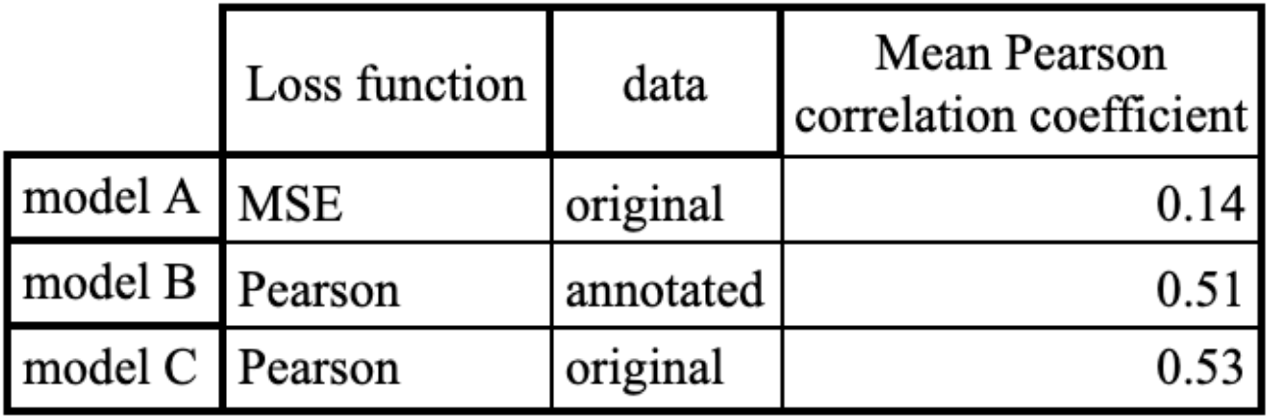
Comparison of Multiple Models.

|  | Loss function | data | Mean Pearson correlation coefficient |
| --- | --- | --- | --- |
| model A | MSE | original | 0.14 |
| model B | Pearson | annotated | 0.51 |
| model C | Pearson | original | 0.53 |

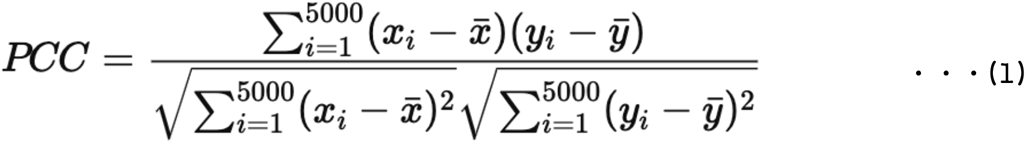

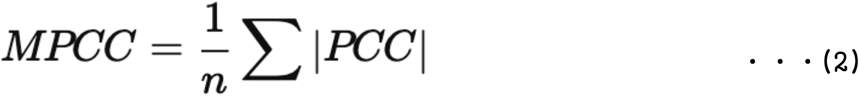

Equations 1 and 2 show the formula for calculating the PCC (Pearson correlation coefficient) within the mini-batch intervals of the patient data, divided by the total number of data points. In this study, we refer to this as MPCC (Mean Pearson correlation coefficient) and use it to evaluate and train the models. Specifically, the data for each subject was divided into 5000-sample segments, and Pearson correlation coefficient was calculated for each segment.

From the comparison of models A and C, we found that the MPCC value was higher by 0.39 points for the model using the Pearson correlation coefficient, indicating better accuracy. Additionally, the comparison of models B and C, where model B used data annotated by physicians, showed that the model trained with the original unannotated data as ground truth (model C) achieved a higher MPCC value and better accuracy.

To verify whether the use of physician-annotated data for the ground truth labels was beneficial, we compared models B and C. The results indicated that model C, which used the original data, demonstrated better accuracy.

Figure 3 provides an example visualizing the output of the best-performing model over a 30-minute measurement period. The top row represents the predicted data inferred from single-lead ECG, the middle row shows the ground truth data from the non-stress test, and the bottom row displays the original single-lead ECG. The MPCC between the ground truth labels and the predicted waveforms in this figure is 0.76. Figure 4 presents multiple visualizations of the evaluation data alongside their corresponding MPCC values.

**Figure 3.**
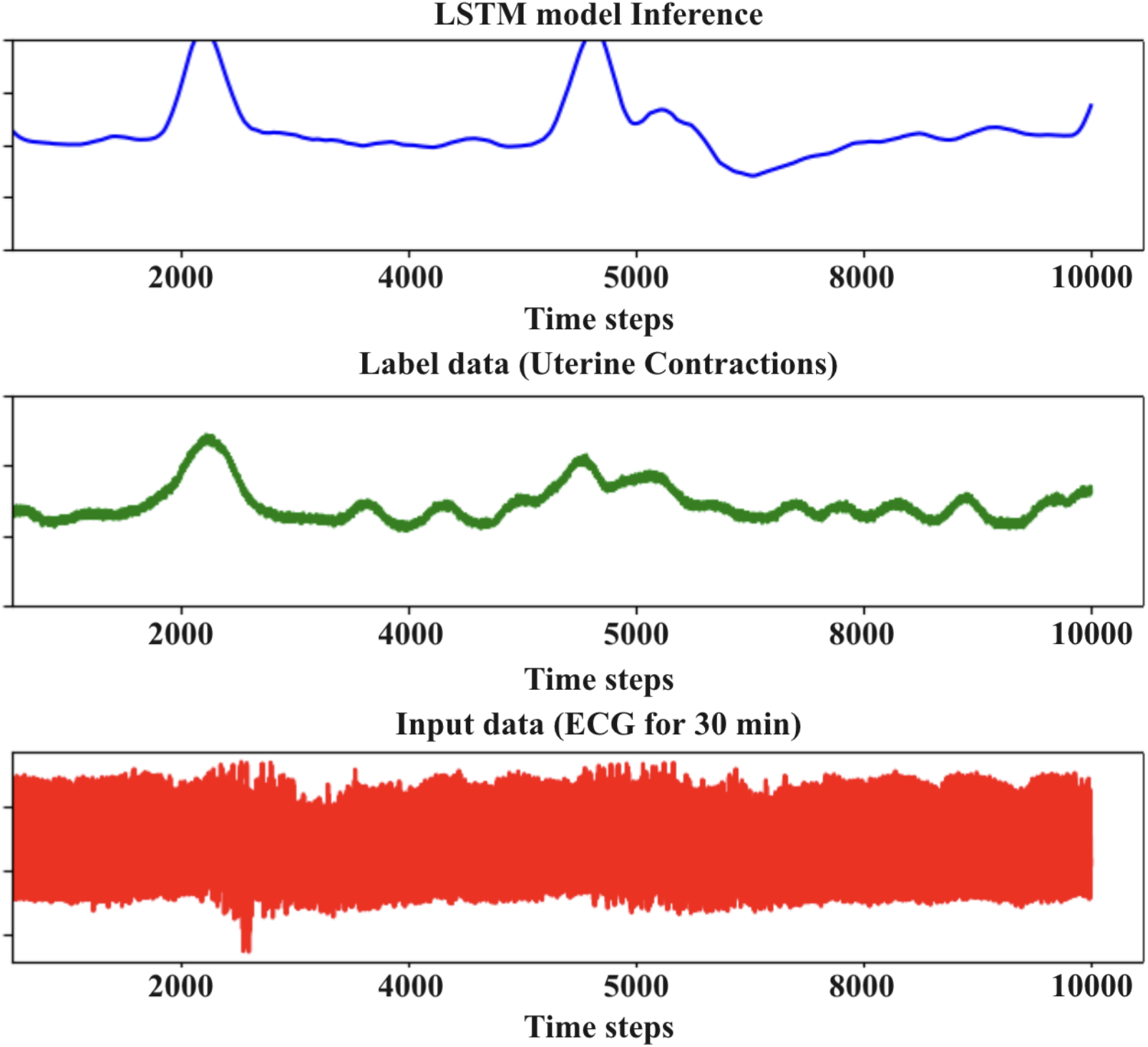
A figure visualizing the evaluation data using the best model.

**Figure 4.**
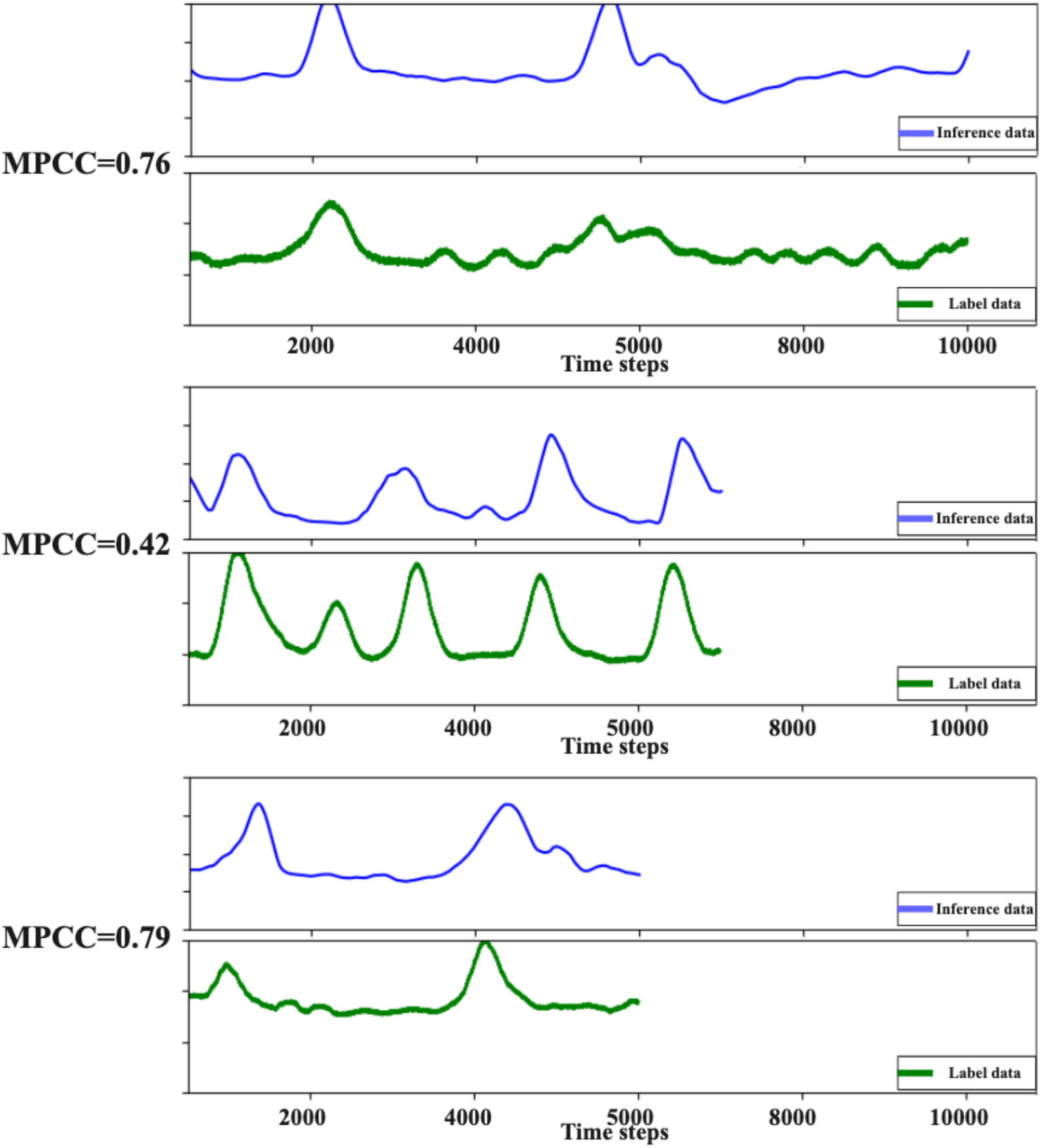
A figure visualizing multiple evaluation data along with the MPCC.

## Discussion

### Medical perspective

We have constructed, for the first time globally, a trained algorithm that accurately infers uterine contraction waveforms from maternal ECGs. Today, ECGs are biological signals that can be easily acquired using wearable devices. Accurate evaluation of uterine contractions outside clinical environments is expected to contribute to the prevention of precipitous labor and, in the future, to the early identification of patients at high risk of preterm birth. On the other hand, the appropriateness of methods for monitoring pregnancy status outside hospital settings (home monitoring) is currently considered to be at the investigational stage (36), and proactive recommendations have been avoided due to the possibility of leading to unnecessary medical consultations (37). However, the cited articles (38)(39) that form the basis of these statements were both published in the 20th century and assume that the interpretation of uterine contractions outside hospital settings is entirely entrusted to mothers with little medical knowledge. Given the remarkable improvements in the inference accuracy of artificial intelligence in recent years, we believe it is necessary to reconsider the utility of home monitoring.

We believe that AI-driven home monitoring will usher in a new era. This is because, in the medical field, there are already numerous cases where AI understands mapping relationships that surpass human cognitive limits and provides valuable information (23)(24)(25)(26)(40). Of course, the approach presented in this study alone is insufficient. For instance, in the prevention of preterm birth, combining our method with conventional treatment protocols for threatened preterm labor could improve pre-test probabilities, reduce unnecessary treatments, and enable effective interventions for high-risk patients.

It is widely known that action potentials occur in association with uterine contractions (41). Techniques to extract these action potentials from the maternal abdomen have been applied as electrohysterograms (or electromyograms) (42)(41). Furthermore, there are known methods for effectively extracting and analyzing uterine contraction– derived action potentials obtained via electrohysterograms (EHG) using deep learning (43). However, electrohysterograms (or electromyograms) assume acquisition from the maternal abdomen, and to our knowledge, there have been no previous cases where action potentials associated with uterine contractions are measured from action potentials obtainable from the arm (i.e.,ECGs), as in our current study. Given that the electrical circuit characteristics of the human body have been known for a long time (44), we considered it highly possible that the electrical activity originating from uterine contractions, detectable from the maternal abdomen, could also be acquired from the arm.

In addition to the electrical signals associated with maternal cardiac contractions detected in ECGs (45), various noises and artifacts such as patient movements, baseline oscillations, electromyographic electrical activity, and electrode movement are also known to be present (46). Detecting uterine contractions from ECGs affected by these noises is considered beyond the limits of human cognition. However, we have demonstrated that the remarkable advancements in deep learning in recent years can overcome this problem. In this study, we have shown that it is possible to effectively infer uterine contraction waveforms from the complex electrical signals included in ECGs obtained from the maternal arm using deep learning. Moreover, several studies have observed that electrohysterography (EHG) shows better results compared to cardiotocography using strain sensors commonly employed in daily clinical practice (47)(41). This suggests that our approach may potentially provide more accurate information about uterine contractions than conventional cardiotocography in the future.

The cardiotocogram (CTG), used as a conventional external pressure gauge, is an indirect indicator that graphs the shifts in resistance values from strain gauges built into the monitor. It is difficult to define a clear correlation between the amplitude of the waveform and intrauterine pressure—the absolute indicator of uterine contractions — which supports our approach. Since this study analyzes the electrical signals derived from uterine contractions on maternal ECGs using deep learning, it becomes possible to evaluate uterine contractions more directly compared to conventional methods. This suggests the possibility of new interpretations regarding uterine contractions that were difficult to achieve with traditional methods. Although the patients targeted in this study did not include those with preterm births (less than 37 weeks of gestation), uterine contractions are considered one of the evaluation indicators for preterm birth risk (2). Therefore, as a future prospect, similar verification in earlier gestational weeks holds the potential to contribute to the assessment of preterm birth risk.

### Technical perspective

We evaluated models using both MSE and Pearson correlation coefficient as loss functions. The model with higher accuracy was the one using Pearson correlation coefficient, suggesting that for this study, it is preferable to use Pearson correlation coefficient as the loss function. For waveform data such as uterine contractions, where the height and interval of the waveforms vary between individuals, Pearson correlation coefficient was found to be the most suitable evaluation metric for this model.

Additionally, in medical data, the measurement duration often varies between samples, and we found that the mini-batch evaluation using Pearson correlation coefficient that we employed was effective in such cases. There was a bias in the average measurement time between the training and evaluation datasets, due to the limited number of measurement samples and insufficient data volume. By splitting the measurement data and evaluating with fixed mini-batches, we were able to ensure unbiased evaluation across all the data, and we confirmed that differences in measurement duration did not affect accuracy.

During the preprocessing stage, expert annotations were applied to the ground truth uterine contraction data to differentiate between contraction and non-contraction periods, as these distinctions were difficult to make. However, the validation results showed that the model using the original, unannotated data achieved higher accuracy. We believe the reason for the reduced accuracy when using the annotated data is that the model might have detected waveform features of uterine contractions that are undetectable by humans. When these were compared to the ground truth labels, the evaluation accuracy worsened.

Based on these findings, we concluded that using the original data leads to better accuracy, as there are subtle contractions and waveform features that are difficult even for physicians to accurately identify, and annotations of such features may suffer from reproducibility issues. Therefore, using the original data, rather than annotated data, results in improved accuracy.

We are also in the process of validating a model that incorporates patient metadata. Although it is generally considered challenging to integrate metadata, which lacks temporal information, into time-series models such as LSTMs, recent research has explored ways to incorporate metadata into deep learning models (27)(48)(49). We believe that with an increase in sample size, this integration can be realized in the future.

In clinical research, in addition to uterine contraction waveforms, we also collect data such as gestational age, weight, BMI, and estimated fetal weight. These factors are known to influence the shape of the ECGs (50), with obesity affecting ECG morphology, the height of the uterine fundus changing with gestational age, and consequent changes in the diaphragm (51). Additionally, variations in maternal circulatory blood flow due to gestational age (52) have been observed. By fusing these factors into the model and considering their effects on the ECGs, we expect to improve the model’s inference accuracy.

In the preprocessing stage, we extracted the peaks of the P-wave, Q-wave, R-wave, S-wave, and baseline wave from the input ECG data before feeding them into the model. This method allowed us to capture the essential characteristics of the ECG while excluding unnecessary components. During the validation phase, we observed that excluding some of these waveform peaks did not significantly affect the prediction accuracy. This suggests the potential to identify which ECG waveform features are influenced by uterine contractions through further validation. By clarifying the relationship between these features, it is likely that a more lightweight model can be developed.

The constructed model achieved an MPCC of 0.53 across the evaluation dataset. This value generally indicates that there is a correlation between the true label and the predicted waveform. The LSTM, which includes a hidden state for short-term memory and a cell state for long-term memory, appears to have functioned effectively. These findings suggest that the characteristics of uterine contraction data may be well-suited for waveform estimation models using LSTM.

In this study, we developed an LSTM model, and while a certain degree of correlation was observed in the model’s predictions, several challenges remain in achieving higher accuracy. LSTM models have difficulty retaining information over extremely long sequences, and when it comes to accurately capturing the periodicity of uterine contractions, the model may reach the limits of its memory retention capabilities.

As we accumulate more data in the future, we plan to explore models with multi-layer structures and self-attention mechanisms, such as Transformer, which generally have more parameters than LSTM. Models that utilize attention mechanisms, such as Transformer, have the potential to model dependencies between all data points and accurately capture important information from distant past events. By applying the insights gained from this study, we believe that we can resolve issues related to optimal preprocessing for large datasets and ultimately achieve higher accuracy.

In particular, pre-training using models based on the Transformer architecture, such as BERT, which is widely used in large language models (LLMs), may contribute to generating outputs closer to uterine contraction waveforms and predicting future contractions.

However, as noted in the cited literature(53), newer models like XLSTM have been proposed for time-series data. Given the characteristics of periodic time-series data, models based on LSTM may still provide more effective learning compared to attention mechanisms in this research context.

## Limitation

Firstly, the sample size of this study was limited to only 57 cases, which raises concerns about the potential risk of overfitting due to the specific characteristics of the population. Although increasing the number of cases in future research could mitigate this issue, in the present study, we addressed the sample size limitation by applying mini-batch learning to the time-series data analysis for each individual and creating datasets based on window lengths. This approach allowed us to generate multiple datasets from the same subjects, thereby enhancing the effective sample size.

Additionally, there is a possibility that electrical signals originating from maternal muscles other than the uterine smooth muscles (e.g., rectus abdominis or biceps brachii) associated with uterine contractions were captured in this study. Future work should aim to clearly distinguish these signals and further investigate the transparency of the AI model to ensure accurate interpretation of uterine contractions.

In clinical practice, awareness of uterine contractions does not always accompany the contractions, and a certain number of pregnant women do not experience conscious sensations of uterine contractions. In such cases, complex electrical signals likely arise from both uterine muscle-derived electrical activity and unconscious maternal muscle-generated electrical signals. The ability of our study to effectively detect uterine contractions through deep learning from these complex electrical signals is therefore highly significant.

Furthermore, another medical aspect related to the elucidation of the deep learning black box is the possibility that the AI may be interpreting biological information other than the complex electrical signals associated with uterine contractions. For example, maternal heart rate is known to change during uterine contractions(54), raising concerns that the algorithm might be capturing merely changes in heart rate rather than the electrical signals themselves. To address this concern, we removed the temporal information of the ECGs during the data preprocessing stage and analyzed the data by treating the time intervals between baseline and PQRST waveforms as fixed intervals. Specifically, we incorporated the ECG waveform information at equal intervals for each LSTM time step. This processing allowed the model to exclude explicit temporal information for each heartbeat, effectively eliminating time-related data from the input. We believe that this approach likely eliminates the possibility that uterine contractions are being inferred solely based on heart rate changes. However, it is also probable that incorporating temporal information could enhance the model’s accuracy, and thus we intend to investigate this in future studies.

## Conclusion

This is the world’s first trial to detect uterine contractions from maternal ECGs using deep learning algorithms. In this pioneering study, we revealed a mapping relationship between the electrical activity associated with uterine contractions and signals obtainable from maternal ECGs. We successfully developed a deep learning algorithm capable of accurately inferring uterine contraction waveforms from maternal ECGs. Our proposed sequence-to-sequence (seq2seq) LSTM model overcomes the limitations of conventional LSTM models in handling long-term time series data such as ECGs and uterine contractions. Although we focused on patients beyond 37 weeks of gestation—differing from those at risk of preterm labor—our findings have the potential to apply to patients before 37 weeks, considering the continuous physiological changes throughout pregnancy. Given that ECGs can be easily obtained using wearable devices, this approach suggests the potential to provide healthcare professionals with objective and accurate information on uterine contractions, a crucial indicator of preterm labor, even in resource-limited settings outside hospital environments.

## Acknowledgements

The authors would like to express their sincere gratitude to all those who contributed to the completion of this study. In particular, they are deeply thankful to the patients for providing invaluable data, as well as to the medical staff at Saintpaulia Misao Ladies Hospital for their essential support. The authors also appreciate ATR-Promotions Inc. for supplying the biosensors used in data collection, and the developers of the Neurokit2 Python library, which was crucial for ECG feature extraction. Finally, the dedicated efforts of the team at nonat Inc. and all other contributors were instrumental in making this research possible. Without the cooperation and assistance of these individuals and organizations, this study would not have been successful.

## References

1. Langen ES, Weiner SJ, Bloom SL, Rouse DJ, Varner MW, Reddy UM, et al. Association of Cervical Effacement With the Rate of Cervical Change in Labor Among Nulliparous Women. Obstet Gynecol. 2016 Mar;127(3):489–95.

2. Haas DM, Imperiale TF, Kirkpatrick PR, Klein RW, Zollinger TW, Golichowski AM. Tocolytic therapy: a meta-analysis and decision analysis. Obstet Gynecol. 2009 Mar;113(3):585–94.

3. Dagklis T, Akolekar R, Villalain C, Tsakiridis I, Kesrouani A, Tekay A, et al. Management of preterm labor: Clinical practice guideline and recommendation by the WAPM-World Association of Perinatal Medicine and the PMF-Perinatal Medicine Foundation. Eur J Obstet Gynecol Reprod Biol. 2023 Dec;291:196–205.

4. Transvaginal cervical length measurement for prediction of preterm birth in women with threatened preterm labor: a meta‐analysis - Sotiriadis - 2010 - Ultrasound in Obstetrics & Gynecology - Wiley Online Library [Internet]. [cited 2024 Sep 28]. Available from: https://obgyn.onlinelibrary.wiley.com/doi/10.1002/uog.7457

5. Born too soon: decade of action on preterm birth [Internet]. [cited 2024 Sep 28]. Available from: https://www.who.int/publications/i/item/9789240073890

6. Epidemiology and causes of preterm birth - PubMed [Internet]. [cited 2024 Sep 28]. Available from: https://pubmed.ncbi.nlm.nih.gov/18177778/

7. Childbearing beyond maternal age 50 and fetal outcomes in the United States - PubMed [Internet]. [cited 2024 Sep 28]. Available from: https://pubmed.ncbi.nlm.nih.gov/14672478/

8. Impact of maternal age on obstetric outcome - PubMed [Internet]. [cited 2024 Sep 28]. Available from: https://pubmed.ncbi.nlm.nih.gov/15863534/

9. Pregnancy outcome at extremely advanced maternal age - PubMed [Internet]. [cited 2024 Sep 28]. Available from: https://pubmed.ncbi.nlm.nih.gov/20965486/

10. Delayed childbearing and its impact on population rate changes in lower birth weight, multiple birth, and preterm delivery - PubMed [Internet]. [cited 2024 Sep 28]. Available from: https://pubmed.ncbi.nlm.nih.gov/11875131/

11. Delayed childbearing and risk of adverse perinatal outcome. A population-based study - PubMed [Internet]. [cited 2024 Sep 28]. Available from: https://pubmed.ncbi.nlm.nih.gov/1640617/

12. Child mortality (under 5 years) [Internet]. [cited 2024 Sep 28]. Available from: https://www.who.int/news-room/fact-sheets/detail/levels-and-trends-in-child-under-5-mortality-in-2020

13. Achana F, Johnson S, Ni Y, Marlow N, Wolke D, Khan K, et al. Economic costs and health utility values associated with extremely preterm birth: Evidence from the EPICure2 cohort study. Paediatr Perinat Epidemiol. 2022 Sep;36(5):696–705.

14. Waitzman NJ, Jalali A, Grosse SD. Preterm birth lifetime costs in the United States in 2016: An update. Semin Perinatol. 2021 Apr;45(3):151390.

15. Chapter 3 The economic case for a shift to prevention [Internet]. [cited 2024 Sep 28]. Available from: https://assets.publishing.service.gov.uk/media/5a7c89deed915d6969f45950/33571_2901304_CMO_Chapter_3.pdf

16. Neonatal Morbidities in Infants Born Late Preterm at 35-36 Weeks of Gestation: A Swedish Nationwide Population-based Study - PubMed [Internet]. [cited 2024 Sep 28]. Available from: https://pubmed.ncbi.nlm.nih.gov/33662344/

17. Effect of late-preterm birth and maternal medical conditions on newborn morbidity risk - PubMed [Internet]. [cited 2024 Sep 28]. Available from: https://pubmed.ncbi.nlm.nih.gov/18245397/

18. Clinical outcomes of near-term infants - PubMed [Internet]. [cited 2024 Sep 28]. Available from: https://pubmed.ncbi.nlm.nih.gov/15286219/

19. Talge NM, Holzman C, Wang J, Lucia V, Gardiner J, Breslau N. Late-preterm birth and its association with cognitive and socioemotional outcomes at 6 years of age. Pediatrics. 2010 Dec;126(6):1124–31.

20. Neonatal morbidity in singleton late preterm infants compared with full-term infants - PubMed [Internet]. [cited 2024 Sep 28]. Available from: https://pubmed.ncbi.nlm.nih.gov/21895764/

21. Asphyxia, Neurologic Morbidity, and Perinatal Mortality in Early-Term and Postterm Birth - PubMed [Internet]. [cited 2024 Sep 28]. Available from: https://pubmed.ncbi.nlm.nih.gov/27235446/

22. Moster D, Lie RT, Markestad T. Long-term medical and social consequences of preterm birth. N Engl J Med. 2008 Jul 17;359(3):262–73.

23. Artificial intelligence using convolutional neural networks for real-time detection of early esophageal neoplasia in Barrett’s esophagus (with video) - PubMed [Internet]. [cited 2024 Sep 28]. Available from: https://pubmed.ncbi.nlm.nih.gov/31930967/

24. Artificial intelligence for digital and computational pathology | Nature Reviews Bioengineering [Internet]. [cited 2024 Sep 28]. Available from: https://www.nature.com/articles/s44222-023-00096-8

25. Screening for cardiac contractile dysfunction using an artificial intelligence–enabled electrocardiogram | Nature Medicine [Internet]. [cited 2024 Sep 28]. Available from: https://www.nature.com/articles/s41591-018-0240-2

26. Huynh E, Hosny A, Guthier C, Bitterman DS, Petit SF, Haas-Kogan DA, et al. Artificial intelligence in radiation oncology. Nat Rev Clin Oncol. 2020 Dec;17(12):771–81.

27. Scalable and accurate deep learning with electronic health records | npj Digital Medicine [Internet]. [cited 2024 Sep 28]. Available from: https://www.nature.com/articles/s41746-018-0029-1

28. Le MD, Singh Rathour V, Truong QS, Mai Q, Brijesh P, Le N. Multi-module Recurrent Convolutional Neural Network with Transformer Encoder for ECG Arrhythmia Classification. In: 2021 IEEE EMBS International Conference on Biomedical and Health Informatics (BHI) [Internet]. Athens, Greece: IEEE; 2021 [cited 2024 Sep 13]. p. 1–5. Available from: https://ieeexplore.ieee.org/document/9508527/

29. Alamatsaz N, Tabatabaei L s, Yazdchi M, Payan H, Alamatsaz N, Nasimi F. arXiv.org. 2022 [cited 2024 Sep 13]. A lightweight hybrid CNN-LSTM model for ECG-based arrhythmia detection. Available from: https://arxiv.org/abs/2209.00988v1

30. Fotiadou E, Sloun RJG van, Laar Joeh van, Vullings R. A dilated inception CNN-LSTM network for fetal heart rate estimation. Physiol Meas. 2021 May;42(4):045007.

31. Asfaw D, Jordanov I, Impey L, Namburete A, Lee R, Georgieva A. Multimodal Deep Learning for Predicting Adverse Birth Outcomes Based on Early Labour Data. Bioengineering. 2023 Jun;10(6):730.

32. Karita S, Ogawa A, Delcroix M, Nakatani T. Forward-Backward Convolutional LSTM for Acoustic Modeling. In: Interspeech 2017 [Internet]. ISCA; 2017 [cited 2024 Sep 13]. p. 1601–5. Available from: https://www.isca-archive.org/interspeech_2017/karita17_interspeech.html

33. Assessment of uterine contractions in labor and delivery - American Journal of Obstetrics & Gynecology [Internet]. [cited 2024 Sep 30]. Available from: https://www.ajog.org/article/S0002-9378(22)00724-4/fulltext

34. Aslami F, Sripian P. Comparative Study of rPPG-Based Methods Performance at the Various Skin Tones. In Japan Society of Kansei Engineering; 2024 [cited 2024 Sep 14]. p. 1–4. Available from: https://www.jstage.jst.go.jp/article/isase/ISASE2024/0/ISASE2024_1_15/_article

35. Li L, Chen C, Pan L, Zhang J, Xiang Y. Video is All You Need: Attacking PPG-based Biometric Authentication [Internet]. arXiv; 2022 [cited 2024 Sep 14]. Available from: http://arxiv.org/abs/2203.00928

36. Urquhart C, Currell R, Harlow F, Callow L. Home uterine monitoring for detecting preterm labour. Cochrane Database Syst Rev. 2017 Feb 15;2017(2):CD006172.

37. Prediction and Prevention of Spontaneous Preterm Birth | ACOG [Internet]. [cited 2024 Sep 28]. Available from: https://www.acog.org/clinical/clinical-guidance/practice-bulletin/articles/2021/08/prediction-and-prevention-of-spontaneous-preterm-birth

38. A multicenter randomized controlled trial of home uterine monitoring: active versus sham device. The Collaborative Home Uterine Monitoring Study (CHUMS) Group. Am J Obstet Gynecol. 1995 Oct;173(4):1120–7.

39. Monitoring Women at Risk for Preterm Labor | New England Journal of Medicine [Internet]. [cited 2024 Sep 28]. Available from: https://www.nejm.org/doi/full/10.1056/NEJM199801013380103

40. Deep Neural Networks Can Predict New-Onset Atrial Fibrillation From the 12-Lead ECG and Help Identify Those at Risk of Atrial Fibrillation–Related Stroke | Circulation [Internet]. [cited 2024 Sep 28]. Available from: https://www.ahajournals.org/doi/10.1161/CIRCULATIONAHA.120.047829

41. Garfield RE, Maner WL. Physiology and Electrical Activity of Uterine Contractions. Semin Cell Dev Biol. 2007 Jun;18(3):289–95.

42. Uterine electromyography: a critical review - PubMed [Internet]. [cited 2024 Sep 28]. Available from: https://pubmed.ncbi.nlm.nih.gov/8267082/

43. Automatic recognition of uterine contractions with electrohysterogram signals based on the zero-crossing rate | Scientific Reports [Internet]. [cited 2024 Sep 28]. Available from: https://www.nature.com/articles/s41598-021-81492-1

44. The human body as an electric circuit - PubMed [Internet]. [cited 2024 Sep 28]. Available from: https://pubmed.ncbi.nlm.nih.gov/15335590/

45. Ashley EA, Niebauer J. Conquering the ECG. In: Cardiology Explained [Internet]. Remedica; 2004 [cited 2024 Sep 28]. Available from: https://www.ncbi.nlm.nih.gov/books/NBK2214/

46. Characterization of noise in long-term ECG monitoring with machine learning based on clinical criteria - PMC [Internet]. [cited 2024 Sep 28]. Available from: https://www.ncbi.nlm.nih.gov/pmc/articles/PMC10412684/

47. R P, S SD. Acquisition and Analysis of Electrohysterogram Signal. J Med Syst. 2020 Feb 10;44(3):66.

48. Stahlschmidt SR, Ulfenborg B, Synnergren J. Multimodal deep learning for biomedical data fusion: a review. Brief Bioinform. 2022 Mar 1;23(2):bbab569.

49. Lee G, Nho K, Kang B, Sohn KA, Kim D. Predicting Alzheimer’s disease progression using multi-modal deep learning approach. Sci Rep. 2019 Feb 13;9(1):1952.

50. The Effect of Obesity on Repolarization and Other ECG Parameters [Internet]. [cited 2024 Sep 28]. Available from: https://www.mdpi.com/2077-0383/13/12/3587

51. LoMauro A, Aliverti A. Respiratory physiology of pregnancy. Breathe. 2015 Dec;11(4):297–301.

52. Soma-Pillay P, Catherine NP, Tolppanen H, Mebazaa A, Tolppanen H, Mebazaa A. Physiological changes in pregnancy. Cardiovasc J Afr. 2016;27(2):89–94.

53. Beck M, Pöppel K, Spanring M, Auer A, Prudnikova O, Kopp M, et al. arXiv.org. 2024 [cited 2024 Sep 26]. xLSTM: Extended Long Short-Term Memory. Available from: https://arxiv.org/abs/2405.04517v1

54. Maternal heart rate patterns in the first and second stages of labor - VAN VEEN - 2012 - Acta Obstetricia et Gynecologica Scandinavica - Wiley Online Library [Internet]. [cited 2024 Sep 28]. Available from: https://obgyn.onlinelibrary.wiley.com/doi/10.1111/j.1600-0412.2012.01371.x

